# An image-based assay to quantify changes in proliferation and viability upon drug treatment in 3D microenvironments

**DOI:** 10.1101/312504

**Authors:** Vasanth S. Murali, Bo-Jui Chang, Reto Fiolka, Gaudenz Danuser, Murat Can Cobanoglu, Erik S. Welf

## Abstract

**Background:** Every biological experiment requires a choice of throughput balanced against physiological relevance. Most primary drugs screens neglect critical parameters such as microenvironmental conditions, cell-cell heterogeneity, and specific readouts of cell fate for the sake of throughput.

**Methods:** Here we describe a methodology to quantify proliferation and viability of single cells in 3D culture conditions by leveraging automated microscopy and image analysis to facilitate reliable and high-throughput measurements. We detail experimental conditions that can be adjusted to increase either throughput or robustness of the assay, and we provide a stand alone image analysis program for users who wish to implement this 3D drug screening assay in high throughput.

**Results:** We demonstrate this approach by evaluating a combination of RAF and MEK inhibitors on melanoma cells, showing that cells cultured in 3D collagen-based matrices are more sensitive than cells grown in 2D culture, and that cell proliferation is much more sensitive than cell viability. We also find that cells grown in 3D cultured spheroids exhibit equivalent sensitivity to single cells grown in 3D collagen, suggesting that for the case of melanoma, a 3D single cell model may be equally effective for drug identification as 3D spheroids models. The single cell resolution of this approach enables stratification of heterogeneous populations of cells into differentially responsive subtypes upon drug treatment, which we demonstrate by determining the effect of RAK/MEK inhibition on melanoma cells co-cultured with fibroblasts. Furthermore, we show that spheroids grown from single cells exhibit dramatic heterogeneity to drug response, suggesting that heritable drug resistance can arise stochastically in single cells but be retained by subsequent generations.

**Conclusion:** In summary, image-based analysis renders cell fate detection robust, sensitive, and high-throughput, enabling cell fate evaluation of single cells in more complex microenvironmental conditions.

## Background

A fundamental paradigm in cell biological studies involves evaluation of cell responses to molecular perturbations. These perturbations typically leverage gene disruptions or pharmaceutical intervention to affect protein abundance or activity. Often, the responses to such perturbations are measured on cells subjected to *in vitro* conditions under the assumption that the observed phenotypes would translate to similar phenotypes *in vivo* [1, 2]. However, emerging evidence suggests that the three-dimensional characteristics of the cell microenvironment, including matrix composition, dimensionality, and stiffness affect cancer progression [3-6] and there it is unknown how the effects of drug perturbation on 2D assays would translate to more complex assays that in principle represent *in vivo* microenvironment more fully. However, these complex assays are often considered too expensive or difficult to implement for pharmaceutical candidate screening [7-9].

An additional concern during drug screening involves differential responses to perturbation due to cell heterogeneity. Despite observations of drug resistance due to inherent or acquired cell heterogeneity in drug responses [10-12] many high-throughput assays do not measure the single cell responses necessary to determine if a partial response is due to cell heterogeneity. For example, a compound could reduce proliferation in 100% of cells or kill 50% of cells while leaving proliferation unaffected in the remaining cells, but assays measure only cell number would not detect these differences. In addition to the inability to distinguish cytostatic from cytotoxic effects, high throughput assays that employ measurements such as cell luminescence or enzymatic conversion activity as proxy measurements for cell number could be affected by differences in cell state that are neither cytostatic or cytotoxic but simply affect activity of the proxy measure [9]. Thus, there is a need for an assay that is capable of increasing microenvironmental complexity while retaining the simplicity necessary for high throughput drug screening studies.

Here, we demonstrate an approach for quantifying cell fates in response to drug treatment that may be implemented on common equipment in a reasonably high throughput fashion without sacrificing cell fate information or the ability to measure single cell responses. We demonstrate such a microscopy-based using a commonly available epifluorescence microscope and we provide analytical software to facilitate widespread adoption of this approach. Using this methodology, we evaluate the effects of drug treatment on melanoma cells cultured in 3D collagen-based matrices. Such 3D collagen matrices represent a common 3D tissue culture platform [13], which has been used to measure changes to cell migration [14, 15], invasion [16], morphology [17], and proliferation [18] as well as to determine the effects of various drug treatments to cancer cells [19]. Cells were treated with varying doses of the current standard of care for mutant BRaf V600E melanoma patients [20], a combination of Dabrafenib (Dab), a BRaf V600E inhibitor, and Trametinib (Tram), a pan MEK inhibitor, a drug combination which we abbreviate as RMIC. Imaging was performed using a Nikon Ti epifluorescence microscope with motorized stage and image analysis was performed by a custom python program as well as the open source Cell Profiler program. We use this assay to show that cytostatic and cytotoxic effects occur at different drug concentrations and that the cell’s microenvironment influences drug effectiveness. We find that cells grown in 3D spheroids exhibit very similar drug responses to single cells in a 3D microenvironment, in contrast to cells grown on 2D plastic plates which are less sensitive than either single cells in 3D collagen or cells in 3D spheroids. Finally, we implement this assay on a system comprised of two different cell types in co-culture to illustrate how different responses to drug perturbations may be measured in heterogeneous cell populations.

## Results

### Procedure to measure the effects of drug treatment on cell proliferation and viability at single cell resolution

When designing an assay to quantify the effects of drug treatments, we identified three critical design criteria: (1) single cell resolution, which enables identification of cell subtypes that respond differentially to treatment, (2) ability to distinguish cytostatic from cytotoxic effects, and (3) ability to measure cell fates in complex 3D microenvironments. We achieve these design goals by imaging and computational analysis of cells marked for proliferation and viability (Figure 1A). Proliferation can be measured precisely *in situ* by measuring DNA synthesis through the 5-ethynyl-2’-deoxyuridine (EdU) incorporation assay. EdU is a nucleoside analog of thymidine which gets incorporated into DNA during the S-phase of the cell cycle and is detected by a copper catalyzed Click iT reaction that forms a covalent bond between the azide of a fluorophore to EdU. Cell viability is quantified by taking the ratio of cells labeled with Hoechst 33342, a general nuclear marker to those labeled with markers of cell death. A generic marker for dead cells, which lack membrane integrity, involves measurement of ethidium homodimer (EtHd) fluorescence in the nucleus. EtHd is a high affinity nucleic acid stain that emits strong red fluorescence only when bound to DNA but is excluded from cells with viable cell membranes. We also considered that EtHd fluorescence would not be detectable in cells in the early stage of apoptosis; therefore, we use extracellular apopxin to label phosphatidylserine (PS), which flips to the outer cell membrane during apoptosis but is otherwise intracellular, to identify cells in early and intermediate states of apoptosis. Images are acquired throughout a thick 3D sample to create a 3D image stack, which is analyzed slice-by-slice and pixel counts representing markers of cell fate are summed throughout the entire sample (Figure 1B). We employ a ratiometric approach for image quantitation to avoid issues with nuclear overlap and polyploidy which would affect cell counting approaches but not ratiometric approaches. All cell fate measures are reported as pixel ratios so that abnormally shaped cells are counted equally by the nuclear normalization signal (Hoechst) and the marker signal. The large pixel size used for imaging and large number of cells imaged also reduce the effect that any small differences in nuclear morphology would have on the end result. However, researchers should always confirm that their nuclear stain and cell fate markers give clear, bright signal and that nuclear shapes are not dramatically different across conditions as this would bias results.

**Figure 1:**
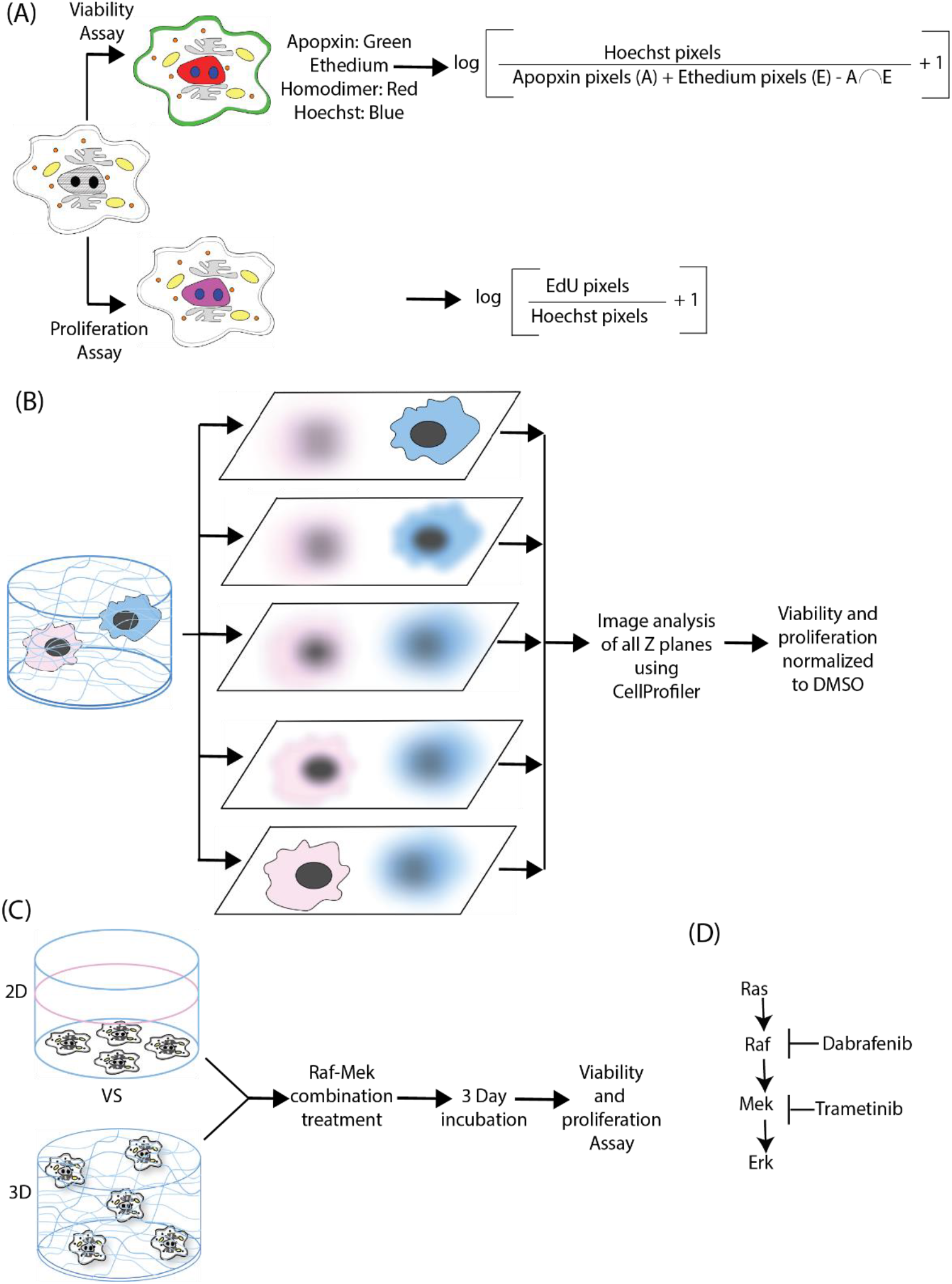
Image-based proliferation and viability measurement of drug-treated cells in 2D and 3D samples. **(A)** Cell viability is measured by image-based quantification of cells positive for extracellular phosphatidyl serine (apopxin) or disrupted plasma membrane (EtHd) and proliferation is measured by image-based quantification of cells positive for EdU incorporation over a 24 hour period. **(B)**. Quantification in 3D samples is performed by obtaining a 3D image stack, analyzing each image slice separately and aggregating data from all image slices. **(C)** Effects of RAF/MEK combination treatment on melanoma cells cultured in standard 2D polystyrene dishes were compared with cells cultured in 3D collagen. (D) RMIC treatment targets mutant BRaf (Dab) Raf and wild type Mek in the MAP kinase pathway.

### Microenvironmental conditions affect melanoma cell sensitivity to RAF/MEK inhibitor combination (RMIC)

We demonstrate the method by quantifying the effects of the RAF/MEK drug combination on mutant BRaf V600E melanoma cells under standard 2D tissue culture and 3D scaffold conditions. We chose collagen I for the 3D scaffold because in addition to being less expensive than other ECM proteins, it is the most abundant ECM protein in the human body [21, 22] and fills the majority of the interstitial spaces in mammals [23, 24], in particular the region of the skin where melanoma originates. ECM compositions that more closely mimic the basement membrane, such as those present in Matrigel, would be more appropriate for cells of epithelial origin. Cells were plated on either 2D polystyrene dishes or embedded in 3D collagen and treated for 72 hours with different concentrations of the RMIC (Figure 1C), which targets the MAP kinase pathway by blocking RAF and MEK (Figure 1D).

After three days of RMIC treatment, cells were assayed for DNA synthesis that occurred during the past 24 hours (i.e. proliferation) and for changes in the viability compared to DMSO-treated controls. As expected, RMIC treatment under both 2D and 3D conditions caused a decrease in proliferation as well as a decrease in the fraction of viable cells (Figure 2).

**Figure 2:**
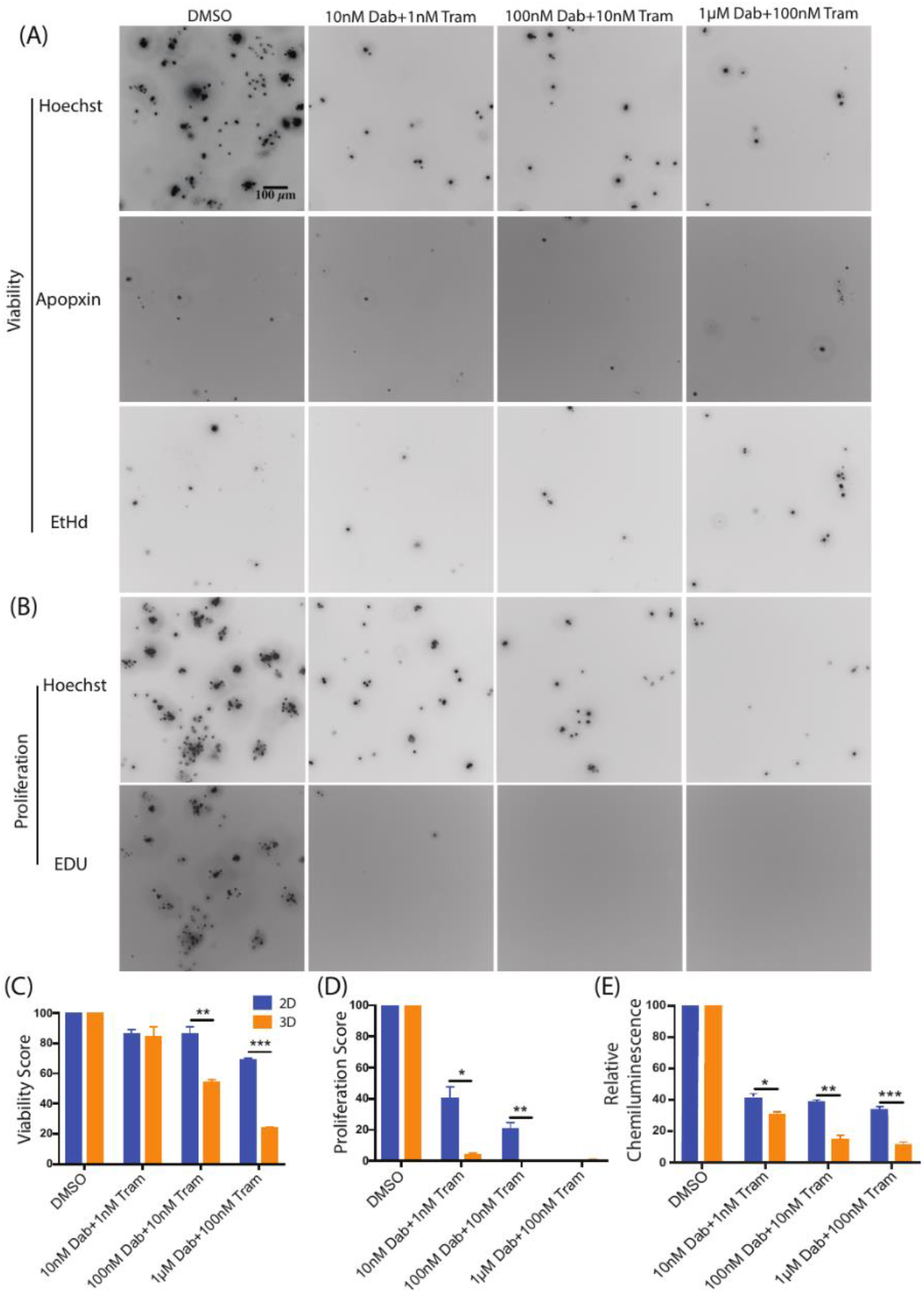
Cell fate assay is capable of identifying differences in proliferation and viability in 2D and 3D microenvironments. Example images show maximum intensity projections of 3D Z-stacks of A375 cells stained with viability markers **(A)** and a proliferation marker **(B)** cultured in 3D collagen and treated with various concentrations of the RAF/MEK inhibitor combination. Plots show aggregate viability score **(C)**, proliferation score **(D)**, and Cell Titer-Glo assay readout **(E)** normalized to DMSO controls, for A375 cells cultured either in 2D dishes or 3D collagen cultures for three independent experimental repeats. Results are shown as average ± SEM (*p < 0.005, ** p < 0.0005, *** p < 0.00005).

However, we note that for most drug concentrations, cells under 3D conditions were significantly more sensitive than cells under 2D conditions. Moreover, the reduction in viability required a RMIC dose 10 times that required to elicit proliferation reduction, suggesting that the RAF/MEK combination elicits more of a cytostatic effect at lower doses. This observation is especially important when we compare these cell fate results to those obtained by bulk cell assays such as the common Cell Titer-Glo assay (Figure 2E). There is an immediate drop off in signal from the Cell Titer-Glo assay, but we are unable to determine if this is due to cell death or a decrease in proliferation. However, the cell fate assay we describe enables identification of the source of the decline in biomass – cell proliferation is rapidly and effectively extinguished even at low RMIC doses, but cell killing only occurs later in time and at higher doses (Supplementary Figure 1). Interestingly, RAF/MEK inhibition is rather ineffective at killing melanoma cells, as A375 cell viability remains between ∼70% and ∼25% under 2D and 3D conditions at the highest concentration of 1µM Dabrafenib + 100nM Trametinib (Figure 2C).

To show that these results are not unique to A375 cells, we performed the assay on melanoma cells derived from two different melanoma patient samples that have been maintained as patient derived xenograft (PDX) models in mice [25]. Cells from both PDX models exhibited proliferation reduction upon RMIC treatment, and cells from both PDX models were more sensitive to proliferation inhibition in 3D conditions than 2D conditions (Figure 3C and 3D).

**Figure 3:**
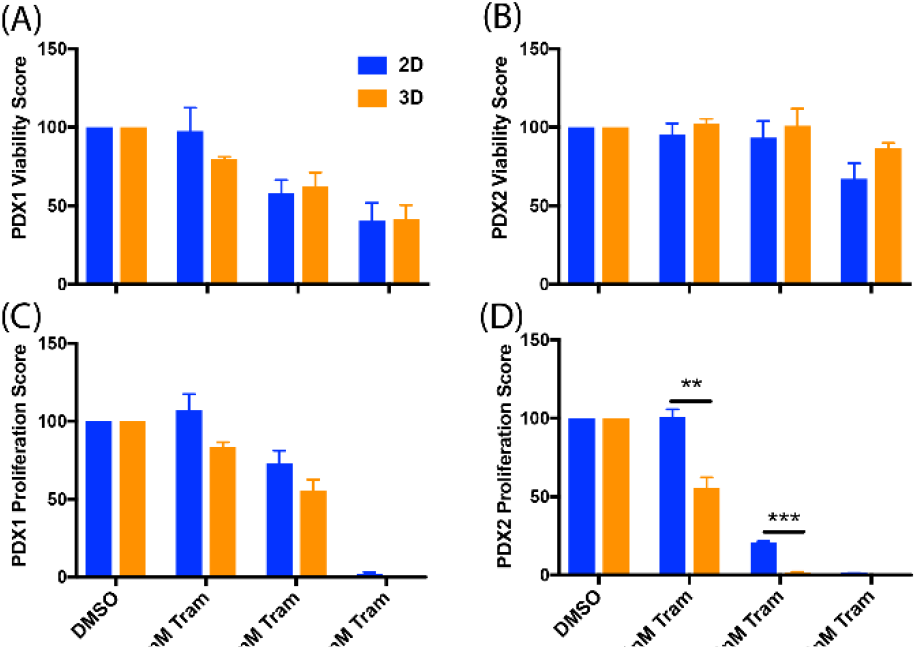
Replication of microenvironmental effects using cells from PDX tumors. Plots show aggregate viability **(A & B)** and proliferation **(C & D)** scores for PDX1, a BRaf V600E mutant melanoma, and PDX2, a BRaf WT melanoma, normalized to DMSO controls, cultured either in 2D dishes or 3D collagen cultures and treated with RMIC treatments as in Figure 2. Data are shown for three independent experimental repeats as average ± SEM (** p < 0.0005, *** p < 0.00005).

The effect on cell viability was more variable, however, as PDX 1 was sensitive to viability reduction regardless of the environmental condition (Figure 3A) but PDX2 exhibited only slight viability reduction at the highest drug concentration (Figure 3B). This result may be explained by the mutational status of PDX2, which expresses wild type (WT) BRaf, and thus is not expected to be sensitive to Dabrafenib, which only targets mutant BRaf. Cells from PDX 2 are expected to remain sensitive to MEK inhibition by Trametinib, however, and this sensitivity explains their reduced proliferation under RMIC.

### Decreased sample size does not cause systematic differences in viability and proliferation in melanoma cells upon RMIC treatment

Although our data shows that cells cultured in 3D microenvironments can respond differently to drug treatment than cells grown on 2D surfaces, a disadvantage of the 3D culture system is the increased experimental cost. Thus, we sought to determine if smaller cell culture volumes may be employed to reduce costs and cell number requirements. We observed no significant difference in the effects of RMIC on viability (Figure 4A) or proliferation (Figure 4B), although greater variability was observed in the 96 well data. This increased variability may be partially attributed to the apparent stiffness associated with the collagen near the walls of the multi-well dish [4, 17, 26], which is known to affect cancer cell fates [4, 27, 28] but would only affect cells close to the wall of the dish. For this assay, we conclude that 24 well or 96 well plates may be used for this assay, giving researchers the option of leveraging the cost savings and increased throughput offered by using 96 well plates at the cost of slightly increased variability compared to larger wells.

**Figure 4:**
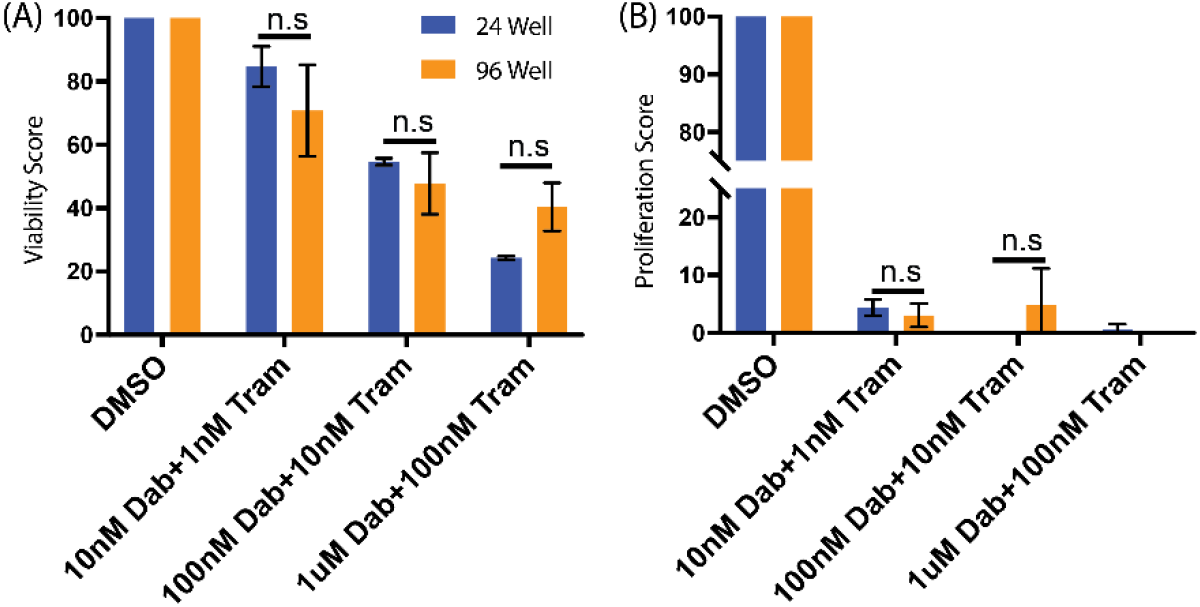
Effect of smaller sample volumes on the image-based assay. A375 melanoma cells cultured in either 24-well or 96-well cell culture dishes were treated with RMIC as a function of concentration as in Figure 2. Plots show viability **(A)** and proliferation **(B)** of the resulting cultures from three independent experimental repeats with results presented as average ± SEM. Statistical comparisons were performed using α = 0.05.

### Flow cytometry is an effective alternative to imaging and computational analysis

Along with microscopy, flow cytometry is an alternative technique for measuring cell fate at single cell resolution, albeit without the spatial context and at the risk of cell extraction and labeling procedures affecting the readout. Here we demonstrate adaptation of the 3D drug treatment assay for flow cytometric measurement by dissociating the collagen gel using collagenase and performing the Click-iT reaction in suspension following collagen digestion according to the manufacturer’s directions (Figure 5A). To discriminate cells positive for EdU from unlabeled cells, a negative control group was assayed, which was not incubated with EdU but was subjected to the Click-iT reaction (Figure 5B). Using this procedure, we find that flow cytometry and image-based analysis result in similar effects on proliferation as a function of RMIC (Figure 5C); however, we do note a reduction in proliferation as measured by the flow cytometry assay compared to the imaging-based assay. Although not statistically significant, this discrepancy may result from reduced sensitivity of the flow cytometer compared to the microscope or to inefficiencies in EdU labeling. Considering this result, we encourage researchers interested in using the flow cytometry approach to consider the potential loss in sensitivity that may result from flow cytometric measurements.

**Figure 5:**
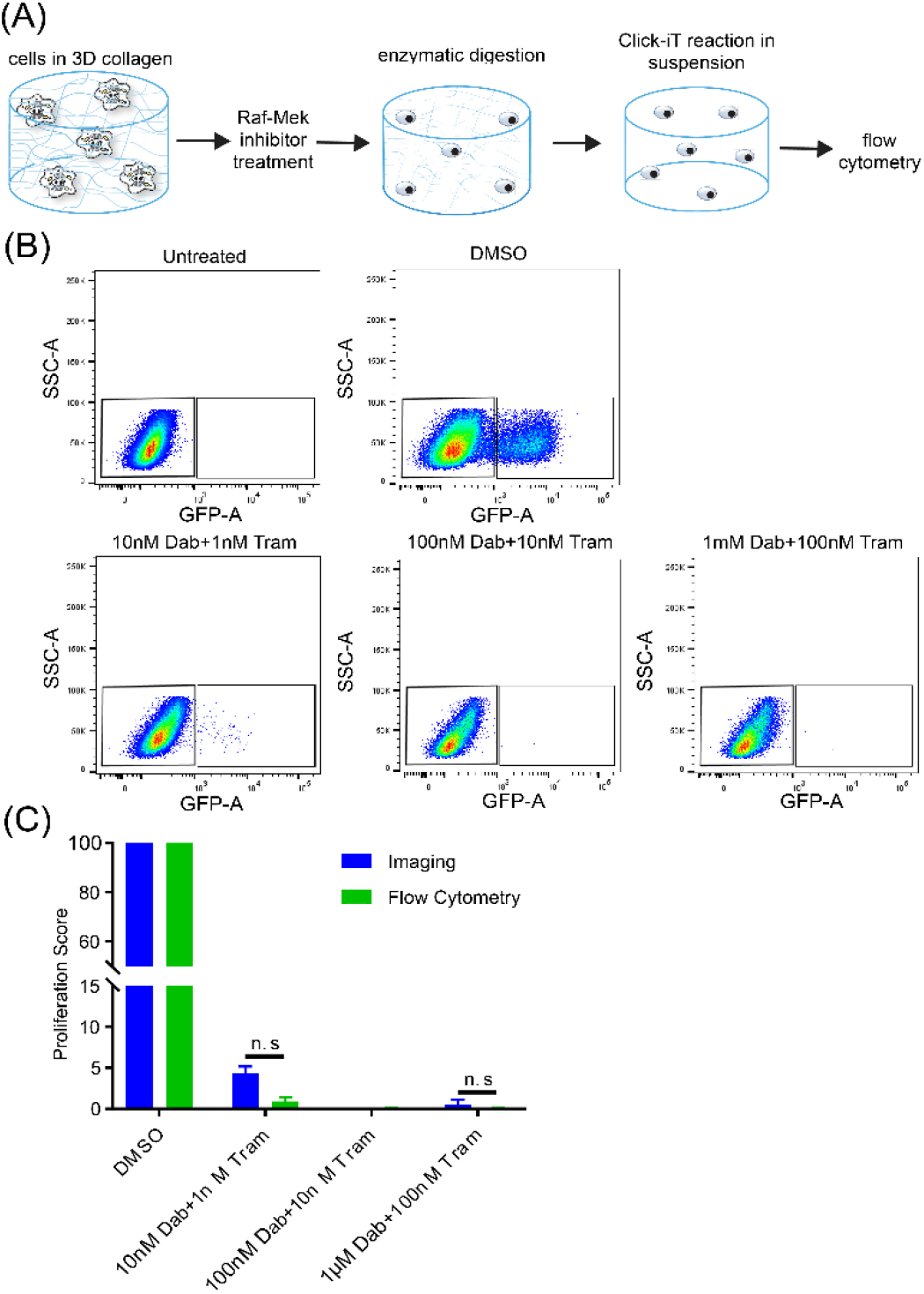
Flow cytometry quantification of proliferation yields similar results to imaging. **(A)** Cells cultured and treated in 3D collagen were released from the collagen prior to fluorescence labeling and flow cytometry. Flow cytometry results were gated based on a negative control **(B)** and the results were compared to imaging platform **(C)**. Plots show data from three independent experimental repeats with results presented as average ± SEM (*** p value < 0.00005).

### Imaging of cells in suspension is an effective alternative to flow cytometry

As an alternative to flow cytometric measurements, we demonstrate the use of image-based analysis for cells cultured in suspension, noting that this measurement procedure may also be used for cells grown in collagen but removed from the collagen prior to measurement. Cells are labeled using the flow cytometry protocol and quantified by placing a drop of cell suspension on a microscope slide and covering the drop with a coverslip (which we term a “coverslip sandwich”) to force all cells into the same image plane (Figure 6A). We demonstrate this approach by measuring the effect of RMIC treatment for cells cultured in suspension, finding that melanoma cells in suspension exhibit similar proliferation inhibition as cells in 3D collagen (Figure 6B and 6C). We also note that this approach enables measurement of cells grown in 3D even if a motorized 3D microscope is not available, rendering the imaging-based 3D assay possible with even basic fluorescence microscopes. The “coverslip sandwich” approach is also particularly useful for measuring samples with low cell numbers, as all the cells in the sample may be concentrated into a single image.

**Figure 6:**
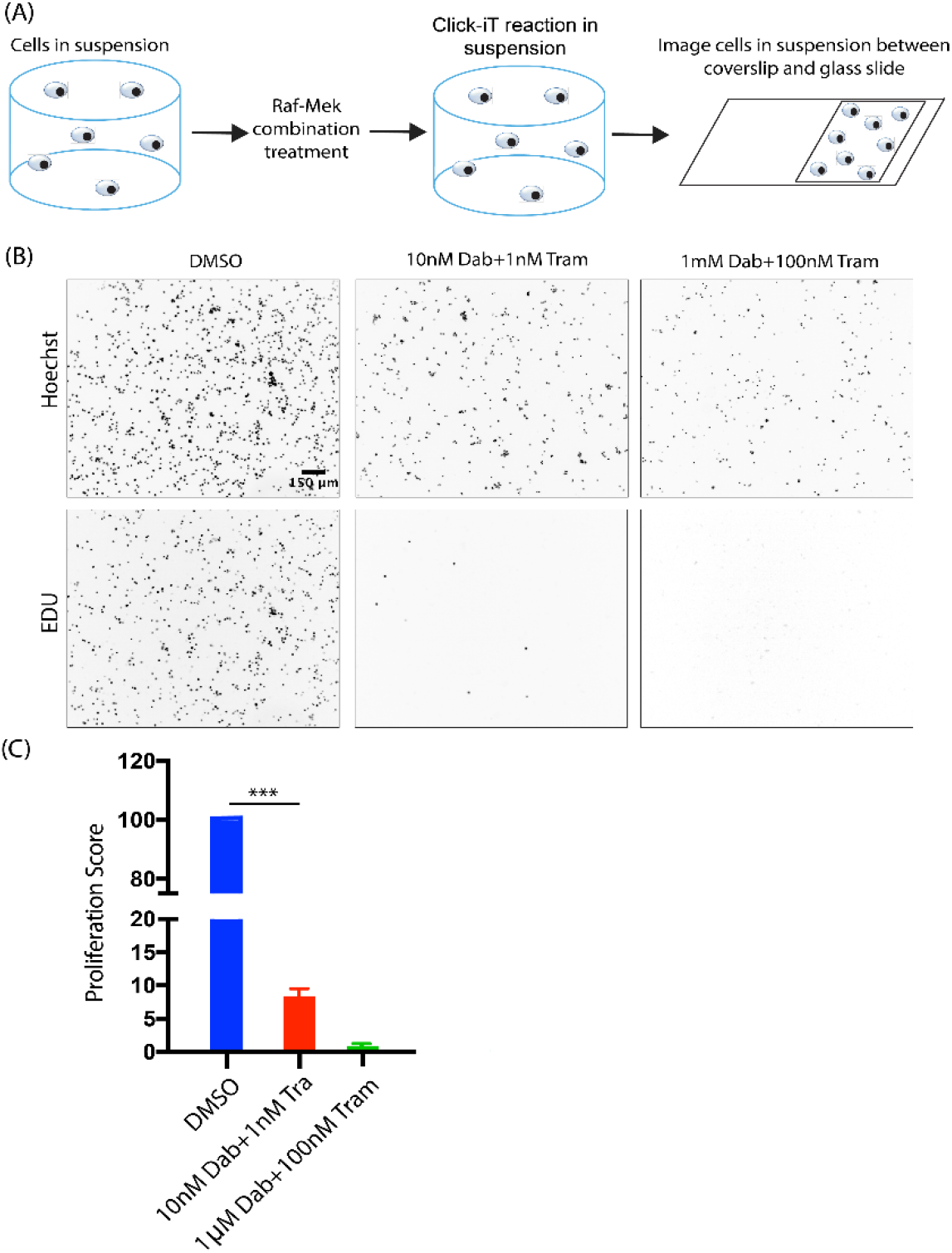
Adaptation of the image-based assay to measure cell fate in suspension. **(A)** Proliferation of A375 cells cultured in suspension was quantified by 2D imaging of cells under a coverslip. **(B)** Example images of suspension cells treated with RMIC acquired as described in **(A)**. **(C)** Aggregate proliferation scores from three independent experimental repeats of cells treated with RMIC in suspension (p value < 0.00005).

### Tumor spheroids exhibit drug sensitivity similar to single cells in 3D

Tumor spheroids are commonly used as 3D cell culture systems as they recapitulate several aspects of *in vivo* tumor morphology [7, 29, 30]. Here we use our cell fate measurement methodology to test the hypothesis that cells in 3D spheroids respond differently to drug treatment than single cells in a 3D microenvironment. Spheroids were formed by seeding single isolated cells in 3D collagen and allowing them to grow into spheroids before RMIC treatment (Figure 7A and Movie 1). Thus, each spheroid is grown from a single cell as it divides but remains adhered to its daughter cells. We employed this approach for generating spheroids because A375 cells formed only loose aggregates of cells via alternative approaches such as the hanging drop method [31, 32].

**Figure 7:**
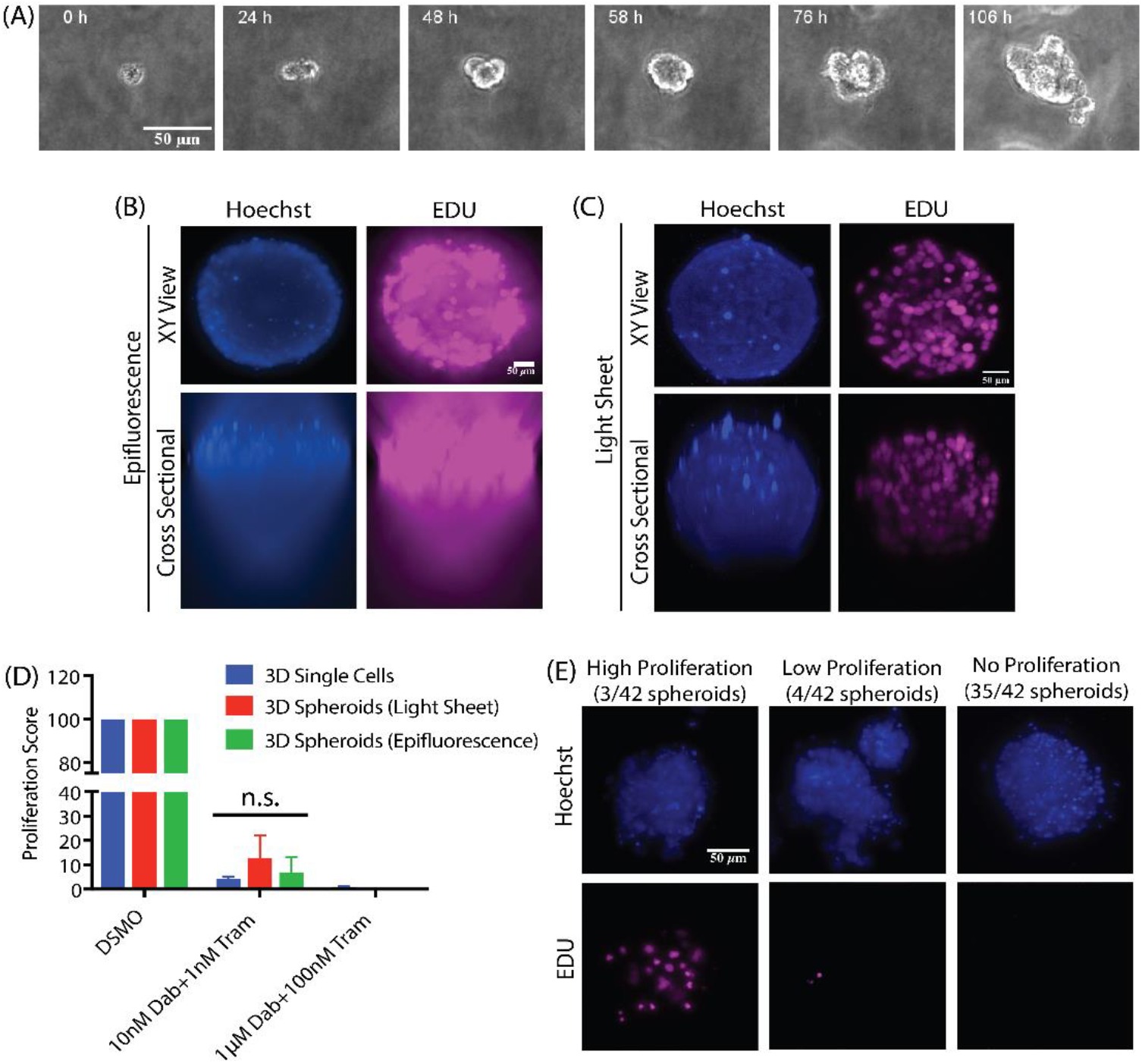
Image-based analysis of RMIC treatment in spheroid models. **(A)** Example images of A375 melanoma spheroids generated in 3D collagen. Example images of spheroids imaged using wide-field epifluorescence **(B)** and LSFM **(C)**. **(D)** Aggregate data comparing the two imaging platforms with similarly treated single cells cultured in 3D collagen from three independent experimental repeats with results presented as average ± SEM, normalized to DMSO control. **(E)** Examples of proliferation heterogeneity in spheroids created from single cells. Of the 42 spheroids imaged, 35 showed no proliferation, 4 spheroids showing low proliferation and 3 spheroids showed high proliferation. Data for the 42 spheroids were a collection from three independent experimental repeats.

A concern that arises from using wide field microscopy to quantify cell fate in such dense samples is that out of focus fluorescence from nearby cells may hinder our ability to identify proliferating cells (Figure 7B). To determine if out of focus fluorescence affects the precision of our measurement, we performed light sheet fluorescence microscopy (LSFM), which limits the amount of out-of-focus light from the 3D sample by exciting fluorescence only in a thin sheet [17, 33]. Compared to wide field imaging, LSFM produced images with individual nuclei that are much easier to distinguish (Figure 7C), improving our ability to identify proliferating cells. Comparison of equivalent samples imaged using LSFM and wide field microscopy demonstrates that the ability to quantify proliferation in spheroids is unaffected by out of focus fluorescence, as samples imaged using LSFM show no significant difference in proliferation compared to samples imaged using epifluorescence microscopy (Figure 7D). More surprisingly, cells treated with RMIC while they are in spheroids exhibited no significant difference in proliferation compared to single cells (Figure 7D).

### RAF/MEK drug resistance in spheroids shows heritability

A benefit of our method for creating melanoma spheroids is that each spheroid is derived from a single cell, allowing us to examine the heritability of drug response. Thus, the striking observation that some spheroids continued proliferating despite RMIC treatment (Figure 7E) while most did not suggests that some heritable drug resistance mechanism of the spheroid initiating cell was passed down to several generations of daughter cells. We also noted that several spheroids possess only one or two proliferative cells, suggesting that drug resistance is either stochastically heritable or transient. The observation that the fraction of proliferating spheroids (3/42) is similar to the fraction of single cells in 3D collagen exhibiting proliferation at the same drug concentration (∼9%) suggests that a similar mechanism of drug resistance may be operating under both single cell and spheroid conditions.

### Ability to discriminate different cell populations reveals that fibroblast co-culture provides little protective effect on melanoma cells treated with RMIC

The ability to distinguish different cell types and identify heterogeneous drug responses is a key feature of our methodology. To demonstrate this capability, we evaluated the effect of RMIC treatment on melanoma cells in 3D co-culture with human foreskin fibroblasts (HFF), which were labeled with an H2B-EGFP marker in order to distinguish them from unlabeled melanoma cells. HFFs are predicted to react differently to RMIC treat because Dabrafenib targets only mutant BRaf and HFFs typically express wild type BRaf. Upon treating HFFs with RMIC we first determined if all GFP positive cells were also Hoechst positive by calculating ratios of GFP to Hoechst pixels (Supplementary Figure 3). We observed more than 80% of the cells to be positive for both GFP and Hoechst pixels irrespective of RMIC treatment. Upon determining viability, they exhibited reduced viability only at high doses of RMIC, (Figure 8C), similar to melanoma cells that express wild type BRaf (Figure 3B). Our image-based assay enabled us to perform an experiment that is impossible using bulk cell assays like CellTiter-Glo, namely to quantify the effect of HFF co-culture on melanoma cells under RMIC treatment while removing the effect of RMIC treatment on HFFs from our measurements (Figure 8). This capability is critical because RMIC treatment affects HFFs at high concentration and this effect would be convoluted with the effect on melanoma cells if a bulk measurement was performed. Indeed, our results show that melanoma viability is reduced at lower RMIC concentrations where HFF viability is unaffected, but that high RMIC affects both HFFs and melanoma cells (Figure 8C). Comparison of the A375 monoculture data with the A375 cells in co-culture reveals little protective effect of fibroblasts in co-culture with melanoma cells, although we do observe a slight reduction in cell killing due to RMIC in melanoma cells at high RMIC concentration.

**Figure 8:**
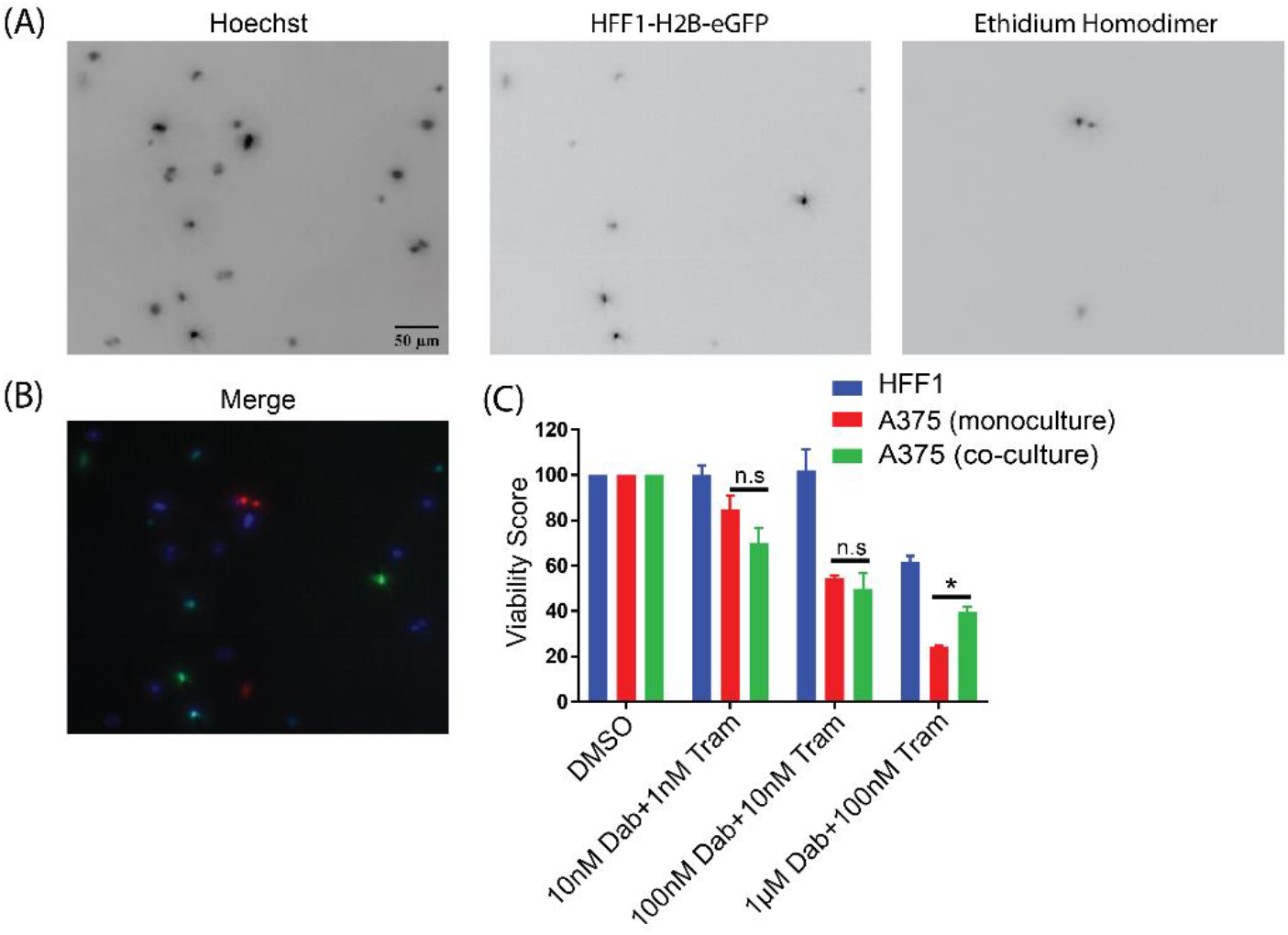
Imaged-based cell fate assay enables measurement of RMIC on melanoma cells in co-culture with fibroblasts. Figure panels in **(A)** show all cells labeled with Hoechst (left), fibroblasts labeled with H2B-eGFP (middle) and dead cells labeled with EtHd (right), with the merged image of all three labels shown in **(B)**. Plot in panel (C) shows the quantification of melanoma and fibroblast viability in co-culture resolved by computational image analysis, compared to melanoma cells in monoculture. Data shown are three independent experimental repeats with results presented as average ± SEM. (* p value < 0.05).

## Discussion

To streamline drug discovery, we must develop methods to evaluate compound efficacy under increasingly complex extracellular microenvironments [34, 35]. However, experimental complexity must be balanced with throughput, cost, reproducibility, and precision. Here, we describe an assay that enables high-throughput compound evaluation without sacrificing single cell resolution, readout specificity, or microenvironmental complexity. Indeed, although we evaluated this assay using 3D collagen scaffolds, it is equally amenable to other scaffolds and would be equally useful for identifying which ECM components affect drug efficacy. We show that wide field 3D fluorescence microscopy is sufficient to quantify cell fate in complex samples including spheroids; however, enzymatic extraction and 2D imaging enables the same level of precision via an even more widely available imaging platform if so desired.

The example results produced via this method also reveal critical insight into how cells respond to drug perturbations in different environments. Indeed, the observation that even a simple 3D collagen microenvironment affects how cells respond to treatment suggests that more careful consideration of the microenvironment may lead to more efficient drug evaluation. For example, the observation that cells in 3D collagen are more sensitive to drug treatment than cells in 2D suggests that many potentially useful compounds might have been ignored because they were tested on cells under 2D conditions. We also show that the standard RAF/MEK treatment is primarily cytostatic and that cytotoxic effects only occur at higher drug concentrations. This result suggests that RAF/MEK treatment may be more effective at slowing tumor growth than eradicating tumor cells and that if slowing tumor growth is the goal then low drug doses may reduce proliferation while reducing side effects compared to higher doses. Similarly, the RAF/MEK treatment affects BRAF wild type melanoma cells and fibroblasts similarly by killing cells only at high concentration, suggesting that the non-targeted effect of the MEK inhibitor Trametinib kills cells indiscriminately and mainly at higher concentrations.

The ability to observe single cell and heritable responses also provides insight into the drug resistance often observed in patients subjected to RAF/MEK inhibitor treatment. For example, if a fraction of melanoma cells are inherently resistant to drug inhibition and the drug effect is primarily cytostatic then it follows that expansion of a drug resistant population will occur. This insight would not be so easily gleaned from conventional methods of evaluating 3D cell culture assays such as chemiluminescence or enzymatic conversion that do not provide cell fate information or single cell resolution [36-38]. Indeed, the effect of RAF/MEK treatment observed in melanoma cells is a combination of proliferation reduction and cell killing that is obscured when measured using Cell Titer-Glo. We also demonstrate the ability to discriminate between different cell populations by testing the hypothesis that fibroblasts in co-culture with melanoma cells can provide a protective effect from RAF/MEK inhibition. The ability to computationally remove fibroblasts from our cell fate measurements, while retaining their effect on melanoma cells, shows that fibroblasts offer little protective effect from RAF/MEK inhibitors for cells co-cultured in 3D collagen. We do observe a small decrease in cell killing at high doses of RMIC, which suggests that the viability effects of RAF/MEK inhibition observed only at high doses may be effected by fibroblasts. We also note that a bulk cell measurement of cells in co-culture would obscure these results -a bulk cell measurement of high RMIC on co-culture would overestimate the effect of RMIC treatment at high doses because it would include the effect of high RMIC doses on fibroblasts.

We also demonstrate how our approach may be used to test the hypothesis that melanoma spheroids offer a more realistic paradigm for drug evaluation than 2D cultures. This is indeed the case – A375 cells grown in spheroids are more sensitive to proliferation inhibition than cells grown on plastic dishes. However, we observe that cells grown as single cells in 3D are equally sensitive to RMIC when compared to cells grown in spheroids. While this phenomenon remains to be evaluated in other systems, the possibility that single cell 3D culture is sufficient to replicate 3D spheroid culture has to potential to greatly reduce experimental costs without information gained from an experiment. Hence, at least for the pathways tested here, drug effects can be tested in the much faster 3D single cell assay and do not require the growth of spheroids. It remains to be determined which condition more faithfully reflects the response of cells to RAF/MEK inhibition *in vivo*.

## Conclusion

The work presented here enables researchers to adopt methodologies for sensitive, informative, high-throughput evaluation of pharmacological perturbations of cells in more realistic microenvironments than standard tissue culture dishes. Our evaluation of different experimental and imaging conditions provides a resource for researcher to make informed decisions regarding experimental design variables such as well size or measurement technique.

## Materials and Methods

### Cell lines

A375 melanoma cells harboring the BRaf V600E mutation were purchased from ATCC (CRL-1619). Two PDX cell models PDX1 and PDX2 were acquired from the University of Michigan via Sean Morrison at UT Southwestern Medical Center. PDX1, like A375, is a BRaf V600E mutant melanoma cell line and PDX2 is a BRaf WT melanoma cell line. HFF-1 was purchased from ATCC (SCRC-1041). In order to discriminate cells in co-culture, the HFF-1 cells were lentivirally transduced with GFP-H2B nuclear marker using the PGK-H2BeGFP system as described by Addgene and selected by flow sorting for GFP positive cells.

### Tissue culture materials

A375 cells were cultured in Dulbecco’s modified eagle media (DMEM), with high glucose and L-glutamine was purchased from Gibco (11965-167), supplemented with 10% fetal bovine serum (FBS), purchased from Thermo Fisher Scientific (11965-167). HFF-1 cells were cultured in DMEM supplemented with 15% FBS. Both PDX1 and PDX2 primary cells were cultured in Dermal Basal Medium purchased from ATCC (PCS-200-030), supplemented with adult melanocyte growth kit purchased from ATCC (PCS-200-042), which includes rh insulin, ascorbic acid, L-Glutamine, epinephrine, CaCl_2_, peptide growth factor and M8 supplement. Phenol red free DMEM with high glucose and L-glutamine for fluorescence imaging was purchased from Thermo Fisher Scientific (21063-045). The phenol red free media was supplemented with 10% FBS. Hanks balanced salt solution (HBSS), with no calcium or magnesium was purchased from Gibco (14170112). PGK-H2BeGFP, lentiviral vector to tag the nucleus of HFF1 cells was purchased from Addgene (Plasmid #21210). Trypsin/EDTA was purchased from Thermo Fisher Scientific (R001100). Trypsin Neutralizer (TN) was purchased from Thermo Fisher Scientific (R-002-100). For 3D culture, bovine collagen I with a molecular weight of ∼300kDa was purchased from Advanced Biomatrix (5005-100). Collagenase Type I powder to breakdown collagen was purchased from Thermo Fisher Scientific (17100017). Dimethyl Sulfoxide (DMSO) was purchased from Thermo Fisher Scientific (BP231-100). Dabrafenib, a BRaf V600-targeting inhibitor (GSK2118436), and Trametinib, a potent MEK1/2 inhibitor (GSK1120212) were purchased from Selleckchem. EtHd, a cell impermeant viability marker, was purchased from Thermo Fisher Scientific (E1169). Apopxin, a phosphatidylserine marker, was purchased from ABCAM (ab176749). Hoechst 33342, nuclear stain was purchased from molecular probes (H3570). Click-iT based imaging kit (C10340) and flow cytometry kit (C10632) were purchased from Thermo Fisher Scientific. CellTiter-Glo was purchased from Promega for 2D (G7571) and 3D (G968A) assays. Tetrafluoroethylene perfluoropropylene (FEP) tubes for spheroid light sheet imaging were purchased from BOLA, Germany (1815-04).

### General cell culture

All 2D cell cultures were performed in 24 well plates. Cells at ∼70% confluency were washed in 1X PBS, trypsinized, and harvested at 500 g for 5 min. The cell pellet was then re-suspended in 1-5 mL media depending on the cell density. The concentration of cells was determined using a cell counter and the dilution was adjusted to 500μL per well for 24 well plates. The cells were seeded at a final density of 15,000 cells/well and incubated overnight to allow attachment to the bottom of the dish following which drug evaluations were performed.

### 3D cultured single cell, co-culture and spheroids

All 3D cell culture experiments were performed in 24 or 96 well plates. Sterile 10X PBS, 1M NaOH and water were pre-warmed in a 37^°C^ water-bath. The multi well plate was pre-warmed in 37^°C^ incubator and an aliquot of 3.2mg/mL collagen was brought to room temperature. *Note*: Cold reagents can lengthen the collagen polymerization time, resulting in cells settling to the bottom of the dish, which does not recapitulate the 3D morphology. To create the cell pellet, cells were trypsinized, harvested and re-suspended in 1-5 mL media depending on the cell density. The concentration of cells was determined using a cell counter and the desired volume of cells were centrifuged to acquire a pellet with the desired cell number. In the case of cells cultured as 3D single cells in 96 well dishes, a seeding density of 30,000 cells/well were used and for 24 well dishes a seeding density of 100,000 cells/well was used. To make spheroids a density of 10,000 cells/well was used. To prepare collagen at a final concentration of 2mg/mL from 3.2mg/mL stock, we combined 100μL 10X PBS, 10μL 1M NaOH, 250μL water and 640μL of 3.2 mg/mL collagen stock. The pH was measured to be between 7-7.4 using pH strips. The cell pellet was re-suspended in 2mg/mL collagen and added to the pre-warmed multi-well plate. The cells were incubated for 30 min in 37^°C^ incubator. Collagen polymerization was observed through change in color of the solution from a transparent solution to a cloudy gel. Media was then added to cells and incubated overnight.

For 3D co-culture experiments HFF1-H2B-eGFP cells were cultured 1:1 with A375 melanoma cells. Briefly, HFF1 mono-culture cells were seeded in 96 well dishes at 20,000 cells/well in a 96 well. We used 20,000 cells because cell numbers higher than 20,000 caused collagen contraction [27]. For the co-culture A375 and HFF1 cells were cultured at 20,000 cells/well with each cell type at 10,000 cells/well. For the co-culture we used both DMSO and HFF1-H2B-eGFP as controls. The cells were setup in 3D like 3D mono-cultures.

A375 melanoma spheroids were created in 3D collagen by suspending single cells in collagen and culturing them over 10 days with frequent replacement of media to form 3D cultured spheroids. For RMIC evaluation in 3D, drug dilution was performed as described below.

### Drug treatment as a function of concentration in 2D and 3D

A375, PDX1, PDX2, HFF1-H2B-eGFP mono-cultures and A375 + HFF1-H2B-eGFP co-cultures were setup in multi-well plates in 3D collagen and 2D culture platforms as described in the methods above. For drug evaluation, a 10mM stock drug concentration was prepared in DMSO. The cells were incubated with three concentrations Dabrafenib and Trametinib: 10nM Dab+1nM Tram, 100nM Dab+10nM Tram and 1μM Dab+100nM Tram with 0.02% (v/v) DMSO as control. A 2X desired concentration of the drug was first prepared in media. This was followed by a 1:1 dilution in media, which yielded the desired final concentration. Controls received the same amount of vehicle (DMSO) as the drug treated cells (0.02% v/v). The cells with drug were then incubated for a total of 72h. After 48h of incubation EdU at a final concentration of 10µM was added. This was done by making 20μM EdU solution which was mixed with 2X drug and subsequently diluted 1:1 with media to give a final concentration of 10μM EdU. This was subsequently incubated in cells for further 24h to make a total drug incubation of 72h.

### Microscopy-based cell fate measurement

Following drug treatment, the media was aspirated from the cells and incubated for 30 minutes with 4μM EtHd, 2μL (per 100μL media) Apopxin and 15μg/mL Hoechst 33342 in fresh phenol red free DMEM supplemented with 10% FBS. Imaging was then performed with a Nikon *Ti* epifluorescence microscope with an OKO temperature and CO_2_ control system regulated at 37°C with 5% CO_2_. The cells in 3D were imaged with a z-step size of 2μm and a total of 251 steps. The filter set for the red channel had an excitation from 540-580nm and emission at 600-660nm, green channel had an excitation from 465-495nm and emission from 515-555nm and blue channel had an excitation at 340-380nm and emission at 435-485nm.

To identify cells undergoing S phase of cell cycle, cells were incubated with modified thymidine analogue EdU for 24 hours and labeled using click chemistry. Following viability imaging after drug treatment, the cells in 2D and 3D were washed with 1X PBS. The cells cultured in 3D were fixed with 4% paraformaldehyde for 30 min at 37^°C^, and the cells in 2D were fixed for 20 min at room temperature. The fixed cells were permeabilized with 0.5% Triton X-100 for 30 min in 3D and 20 min in 2D. *Note:* that the 0.5% Triton X-100 was prepared from a 25% stock solution. The 25% stock should be stored at 4^°C^ and diluted to 0.5% before use. The Click-iT reaction was then performed using the Click-iT EdU assay imaging kit with Alexa Fluor 647 fluorophore. The Click-iT reaction was partially modified from the manufacturer’s protocol such that the Alexa Fluor 647 azide was first diluted 1:100 in DMSO before adding the volume specified by the manufacturer’s protocol, resulting in a concentration 1/100^th^ of the recommended concentration. This dilution needs to be optimized separately for each cell type. If the Alexa Fluor 647 is used directly as per manufacturer’s protocol in 3D the dye will non-specifically bind to the collagen, creating a high background.

### RMIC treatment as a function of concentration on co-cultures

To evaluate identification of heterogeneous cell population using the image based assay we evaluated a co-culture based system. HFF1 cells tagged with H2B-eGFP were co-cultured with un-tagged wild type A375 melanoma cells. The cells in co-culture were seeded at 10,000 cells of each cell type per well in 3D. A control of HFF1-H2B-eGFP cells alone were also setup in 3D. The cells were treated with RMIC as a function of concentration as described above and incubated for a period of 3 days. Following RMIC incubation a modified viability assay was performed. Since the HFF1 cells were tagged with H2B-eGFP, we did not use Apopxin. Viability assays were performed using Hoechst and EtHd.

A control experiment was performed to identify H2B-eGFP pixels intersecting with Hoechst pixels, for HFF1 cells. This would inform how efficiently we can identify HFF1-H2BeGFP cells. Upon imaging and quantification a ratio of eGFP pixels to Hoechst pixels was then quantified for HFF1 cells upon RMIC treatment (Supplementary Figure 2). Viability assay was then performed to determine the positive pixels for Hoechst, eGFP for HFF1 cells and EtHd for total dead cells. The intersection between the positive pixels for each channel were also acquired to identify cells that were both GFP and EtHd positive. This indicates death of HFF1 cells alone. The EtHd pixels that do not overlap with eGFP channel will indicate A375 melanoma cell death. This will give us information on the effect of RMIC on a co-culture based platform.

### Image analysis using Cell Profiler and data processing

To analyze all the 2D and 3D images it was important to identify software that can be used with minimal computational experience. We identified Cell Profiler as an ideal candidate (Carpenter, et al., 2006; Kamentsky, et al., 2011) (http://cellprofiler.org). We used Cell Profiler V 2.2. To operate Cell Profiler we established image analysis pipelines that can accept our multi-point z-stack images and analyzed data for positive pixels from each channel.

The z-stack images were acquired as individual TIFF files. A distinguishing parameter such as a unique name for each channel was used while acquiring 3D and 2D images. In the input module under “NamesAndTypes” we defined the distinguishing name for each channel to help the software separate images from different channels. We used a robust background thresholding method to distinguish foreground (i.e. marked cells) and background regions. The pixel areas of foreground regions were measured for each channel.

While using two dead cell markers (apopxin and EtHd) to identify different phases of cell death, we observed some cells that were marked by both dead cell markers. To identify cells that were counted twice we created a mask to identify overlapping regions and then subtract that area so they are counted only once (Supplementary Figure 3). For analyzing the cell proliferation data the pipeline was modified to two channels and the mask function was removed. Both the viability and proliferation data was exported to a spreadsheet for analysis.

To analyze the viability data we first subtracted the overlapping pixel area of the dead cell channels (apopxin and EtHd) from the total area of the dead cells channels. A log (ratio+1) of the total pixels from hoechst to the dead cell pixels was calculated and averaged over all images acquired for a particular treatment. The average values for each treatment were normalized to the DMSO control; and these values were referred to as viability scores. Analogously, proliferation scores were computed based on averaging and normalization of the log (ratio+1) of the total pixels from Hoechst to EdU pixels.

### Parallel processing image analysis using a custom python program to increase throughput of image analysis

Three challenges with Cell Profiler were (a) the throughput of image analysis upon introducing signal correction functions like illumination correction (b) the absence of automated thresholding and (c) extended processing time. Illumination correction is essential as raw images might have a strong bias towards high intensity values towards the center of the image. To correct for these illumination effects, we first use a local Gaussian with a large variance to smooth the image. We then subtract the background from the foreground and set any resultant negative values to zero because negative intensity is impossible.

To fully automate the image processing, we need to automatically distinguish between background and foreground, hence we need automatic thresholding. To achieve this, we need to tackle two challenges: halo effects from out-of-focus cells, and the absence of biological signal. To deal with halo effects, we adopt a two-pronged strategy. First, we use the thresholding algorithm developed by Yen and colleagues (IEEE Transactions on Image Processing, 1995). We chose this specific algorithm because it seeks to minimize the complexity of the thresholded image, in addition to the standard concerns about seeking a distinct signal level. The assumption of simplicity in the thresholded image makes sense in our domain because the stained biological specimens are all likely to be highly similar and uniform. As such, we observed that the resultant threshold values capture only the in-focus nuclei and not the irregular shapes caused by out-of-focus halo effects. Furthermore, to avoid the irregularities due to halo effects in the extremes of the z-stacks, we only use the images at the center of the z-stack to identify threshold values. Finally, to tackle images devoid of any meaningful biological signal, we compare the threshold value acquired in each image to the mean of the background in the same image. If the threshold is less than ten-fold intense when compared to the background, then we posit that the signal-to-noise ratio is weak due to the absence of any meaningful biological signal.

Finally, the image processing code needs to scale in order to facilitate drug screening. With this goal in mind, we built a highly modular and parallelized solution. In one standard compute node of our cluster, we can process 1,800 images under 3 minutes. By submitting multiple parallel jobs to different nodes, we can process different wells or plates in parallel. When we leverage a high performance computing cluster to run 16 such jobs concurrently, we achieve a throughput of about 10,000 images per minute.

We compared the output of the python script with the output of the CellProfiler pipeline to determine if any systematic differences result from the different analysis approaches. Viability images of A375, PDX1 and PDX2 were analyzed by both approaches and results are compared in (Supplementary Figure 4). The data indicated similar trends to Cell Profiler at lower concentrations for all the three cell lines. However, at the highest concentration of 1µM Dab + 100nM Tram some difference could be observed for A375 and PDX2 models. A possible reason for this could be the illumination correction function that has been used in the Python script to remove background illumination light.

### Evaluating effect of well size by comparing viability and proliferation in 24 and 96 well plates

To understand if well diameter results in variable drug efficacy, we evaluated the effect of RMIC on viability and proliferation as a function of concentration in A375 cells in 24 well and 96 well plates. A375 cells were plated in 2 mg/mL collagen in 3D. The cells were then treated with RMIC as a function of concentration for 3 days. On day 2 of RMIC treatment, 10μM EdU was added for an additional 24h. After 3 days total incubation, viability and proliferation assays were performed as described above.

### Comparing imaging and flow cytometry for RMIC as a function of concentration

To validate the robustness of the 3D imaging platform, we measured RMIC effect on proliferation using the flow cytometry as an independent method. A375 cells were incubated with drugs as a function of concentration. The cells were incubated with drugs for 3 days and on day 2 of treatment, EdU at a final concentration of 10μM was added to the cells. Following incubation with RMIC and EDU, the collagen was washed with Hanks balanced salt solution (HBSS). The collagen was then treated with collagenase as per manufacturer’s protocol for 30 min at 37^°C^ to breakdown the collagen completely. Media with 10% FBS was then added to neutralize the effect of collagenase. A flow cytometry based Click-iT reaction was performed as per manufacturer’s protocol. In brief, the cells were centrifuged to collect the pellet. The pellet was resuspended in 4% PFA for fixation and permeabilized with 1X Tween-20. The Click-iT reaction was performed, and the flow cytometry was gated using a negative control where the cells were not treated with EdU. To analyze the results, a uniform gating was setup by using the FlowJo software with a negative control consisting of cells not subjected to the Click-iT reaction and thus exhibiting only autofluorescence. The treated samples were the gated under the same settings.

#### Comparing imaging and CellTiter Glo for RMIC as a function of concentration

A375 cells were cultured in black walled, glass bottom 96 well dishes. Cells for 2D were seeded at 5,000 cells/well and for 3D at 30,000 cells/well. The RMIC was incubated as a function of concentration as described above for a period of 3 days. After 3 day incubation the cell were treated with the standard CellTiter Glo for 2D and CellTiter Glo-3D reagent for the cells cultured in 3D. The cells in 2D were after addition of reagents were rocked for 2 min and further incubated for 10 min following which the plate was read using a BioTek plate reader. The cells cultured in 3D was incubated with 3D-CellTiter Glo were rocked for 10 min and incubated for a further of 20 min and read.

### Imaging cells in suspension using the Click-iT flow cytometry kit

Since the Click-iT EdU based imaging kit was designed for adherent cells, we aimed to design an imaging based method to determine proliferation for cells in suspension. We thus tested the application to imaging of a flow cytometry kit that is designed for suspended cells. To do so we cultured A375 melanoma cells in ultra low adhesion 6 well plates to keep cells in suspension. The cells were then treated with RMIC and EdU as described above. The Click-iT reaction was performed with the flow cytometry kit as per manufacturer’s protocol and cells were also stained with Hoechst. A drop of the cell suspension after the Click-iT reaction and Hoechst treatment was put on a glass slide and placed under a glass coverslip to, after which images were acquired on the Nikon *Ti* microscope. The cell images for nuclei Hoechst staining and EDU proliferation were analyzed on Cell Profiler.

### Melanoma spheroid preparation for light sheet fluorescence microscopy (LSFM)

Tumor cell spheroids were created from single melanoma cells by culturing cells at low concentration and allowing them to grow into large clusters. Sample holders for light sheet imaging were created as follows. FEP tubes were cut into small pieces with holes made along the walls using a 27G needle in order to allow media exchange across the tube. The tubes were cleaned with 30% bleach for 30 minutes and placed in 100% EtOH overnight. The sterile tubes were dried and pre-warmed in a 37^°C^ incubator. A375 melanoma cells were prepared in 3D collagen as described above and seeded at low density within the FEP tubes. The collagen was allowed to polymerize at 37^°C^, fresh media was added, and then the cells were incubated for 10 days with regular replacement of media. The spheroids then underwent RMIC treatment for 3 days with 24h EdU incubation. Following RMIC treatment and EdU incubation, the Click-iT imaging assay was performed, and the spheroids were imaged on the LSFM.

To image spheroids, a low magnification LSFM was constructed with a large field of view. To increase the field of view, and improve optical penetration, a conventional dual illumination LSFM was built that permits scanning of the beam in the Z-dimension as well as pivoting the light-sheet in the sample plane to reduce shadowing and stripe artifacts [30]. For illumination, four lasers (405nm, 488nm, 561nm, and 640nm, Coherent, OBIS) are combined with dichroic mirrors (MUX Series, Semrock), spatially filtered and expanded with a telescope consisting of an achromatic convex lens (f=50 mm, Thorlabs, AC254-050-A-ML), a pinhole (100 µm, Thorlabs, P100H), and an achromatic convex lens (f=400 mm, Thorlabs, AC254-400-A). An achromatic Galilean beam expander (Thorlabs, GBE02-A) further increases the laser diameter by 2x. All solid-state lasers are directly modulated with analog signals originating from a field-programmable gate array (PCIe-7252R, National Instruments) that have been conditioned with a scaling amplifier (SIM983 and SIM900, Stanford Research Systems).

For light-sheet generation, a cylindrical lens (f=50 mm, Thorlabs, ACY254-050-A) is used to focus the laser illumination into a sheet, which is relayed to the illumination objective (Nikon 10X Plan Fluorite, NA 0.3) with two mirror galvanometers (Thorlabs, GVS001), two coupling lenses (f=50 mm, Thorlabs, AC254-050-A), and a tube lens (f=200 mm, Thorlabs, ITL-200). The Z-galvanometer is conjugate to the back pupil of the illumination objective, whereas the pivot galvanometer is conjugate to the sample. To generate the second illumination arm of the microscope, a polarizing beam splitter (Newport, 10FC16PB.3) is placed after the second coupling lens, and two lenses (f=200 mm, Thorlabs, AC508-200-A) relay the scanned illumination to a second tube lens and illumination objective. Because the counter-propagating light-sheets are orthogonally polarized, there is no interference from the two light-sheets, and a half-wave plate (AHWP10M-600) placed in front of the polarizing beam splitters is used to adjust the intensity of the two light-sheets. To control the light-sheet thickness, a variable slit was placed in the back-pupil plane of the cylindrical lens, which is conjugated to the back-pupil planes of both illumination objectives.

For detection, a 16x, NA 0.8 objective lens (Nikon CFI75 LWD 16X W) and a tube lens (f=200 mm, Edmund optics, 58-520) form the image on a sCMOS camera (Hamamatsu, C11440, ORCA-Flash4.0). A laser line filter (Chroma, ZET405/488/561/640) is placed after the detection objective lens and a filter wheel (Sutter Instrument, Lambda 10-B) equipped with multiple bandpass filters is placed between the tube lens and the camera. The detection objective lens is mounted on a piezo-driven stage (Mad-City Labs, Nano-F450) that provides 450 µm travel range. The sample stage is a combination of a three-axis motorized stage (Sutter Instrument, MP285) and a rotation stage (Physik Instrumente, U-651.03). The microscope is controlled by custom-written LabVIEW software (Coleman Technologies, National Instruments).

#### 3D co-culture experimental procedures and image analysis

To measure the effects of RMIC treatment on fibroblasts alone, we first test the GFP signal detection in fibroblast monoculture. The cells were treated with RMIC as a function of concentration for 3 days. Following treatment, the cells were imaged for viability with Hoechst and EtHd. Apopxin wasn’t used because the HFF-1 cells were GFP tagged. Since it was a mono-culture of only HFF-1 cells tagged with GFP and stained with Hoechst we first tested if all the Hoechst positive cells were also GFP positive. For this we determined the ratio of GFP to Hoechst pixels. Upon quantifying ∼80% showed positivity for GFP and Hoechst pixels irrespective of RMIC concentration (Figure 8A). When quantified for viability HFF-1-H2B-eGFP cells a 100% viability up to 100nM Dab +m 10nM Tram. But at the highest concentration of 1µM Dab + 100nM Tram ∼35% reduction in viability was observed (Figure 8B). A reason for this could be the use of Trametinib which is a pan-MEK inhibitor.

To remove the effect of fibroblasts from our measures of cell fate, the H2B-eGFP pixels were subtracted from the Hoechst pixels. Since the A375 were only stained with Hoechst and not eGFP the subtraction would give us A375 pixels. Likewise, EtHd positive A375 melanoma cells were identified by computationally removing fibroblast cells from the EtHd count. To identify EtHd positive fibroblasts a mask was made to determine the intersection between GFP and EtHd channels and that intersection was then subtracted from the total EtHd positive pixels. Ratios of the EtHd positive A375 cells to Hoechst positive A375 cells were then acquired and normalized to DMSO to give viability results in the co-culture platform.

### Statistical tests

All statistical measurements were performed using Holm-Sidak method, with alpha = 0.05. Each row was analyzed individually without assuming a consistent SD. Results are displayed using p-value of < 0.05: *, < 0.005: ** and <0.0005: ***.

## Declarations

### Abbreviations

Dab: Dabrafenib
Tram: Trametinib
EdU: 5-ethynyl-2’-deoxyuridine
EtHd: ethidium homodimer
PS: phosphatidylserine
RMIC: RAF/MEK inhibitor combination
PDX: patient derived xenograft
WT: wild type
LSFM: light sheet fluorescence microscopy
DMEM: Dulbecco’s modified eagle media
FBS: fetal bovine serum
TN: Trypsin Neutralizer
DMSO: Dimethyl Sulfoxide
FEP: Tetrafluoroethylene perfluoropropylene

### Ethics approval and consent to participate

Not applicable

### Consent for publication

Not applicable

### Availability of data and material

The Python script for high throughput image analysis of cell fate images is available at: www.utsouthwestern.edu/labs/danuser/software/

### Authors’ contributions

V.S.M. and E.S.W. conceived the project and designed the experiments. G.D. and M.C.C. provided intellectual input. M.C.C. wrote the Python image analysis program. V.S.M. and E.S.W. made the figures. V.S.M. performed the experiments. B.J.C. and R.F. constructed the LSFM. V.S.M., B.J.C., R.F., G.D., M.C.C., and E.S.W. wrote the manuscript.

## Acknowledgements

We would like to thank Andrew D. Jamieson for help in building Cell Profiler pipelines, Heather Grossman for cell sorting, Sean Morrison for PDX-derived cell models, Xuexia Jiang for help with file conversions and Meghan Driscoll, Etai Sapoznik and Maralice Conacci-Sorrell for providing valuable feedback on the manuscript.

## Movie 1. Growth of melanoma spheroids from single melanoma cells in 3D collagen

**Supplementary Figure 1:**
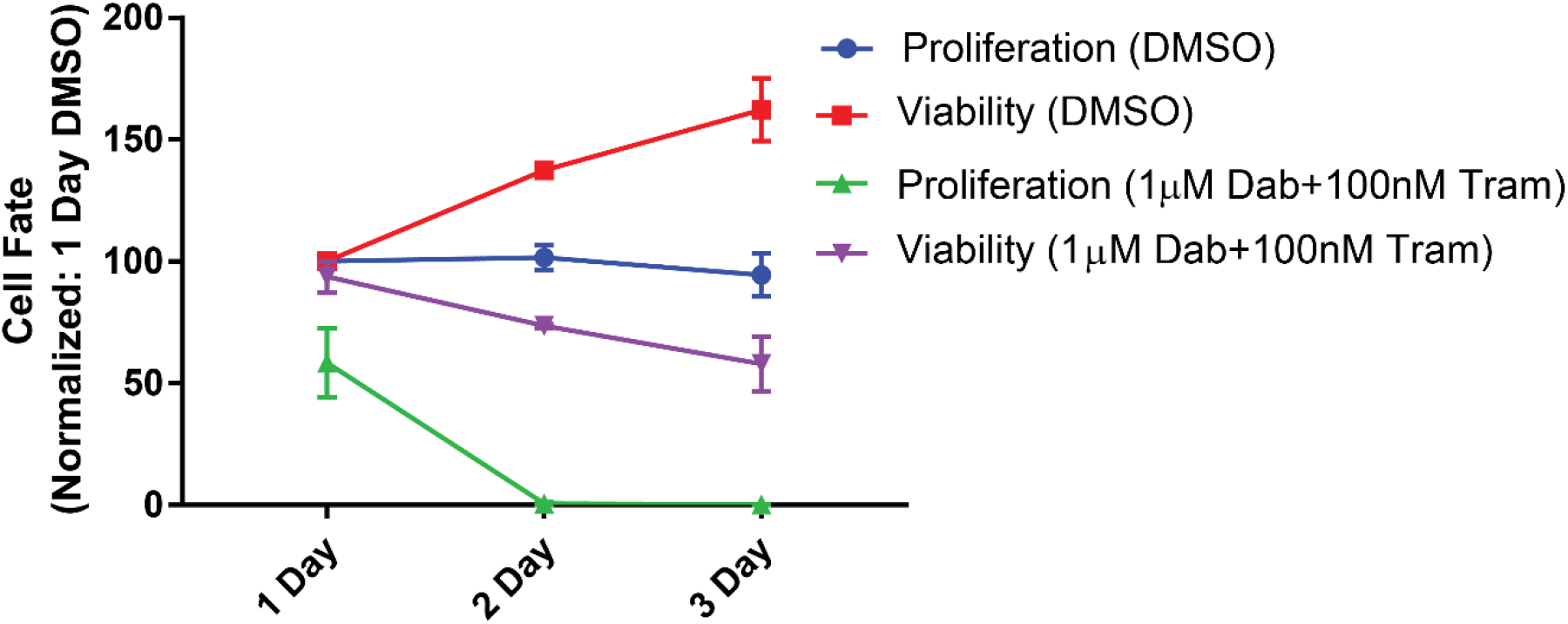
Time course of the effect of RMIC on proliferation. A375 cells treated with RMIC at a concentration of 1µM Dab + 100nM Tram in 3D were incubated for 1, 2 and 3 days.

**Supplementary Figure 2:**
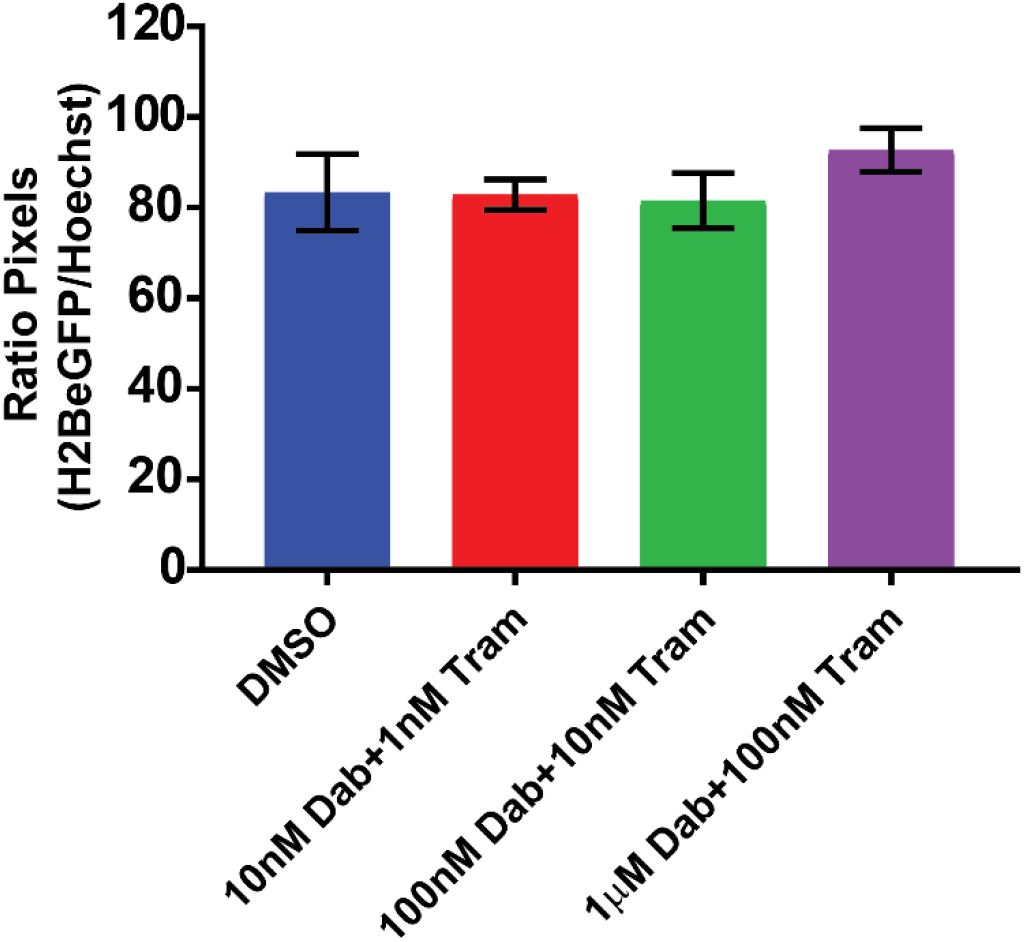
Validation of the image based assay for identifying fibroblasts in 3D culture. HFF1-H2BeGFP cells in 3D collagen were treated with RMIC as a function of concentration and pixels positive for GFP were compared to pixels positive for Hoechst to determine the overlap between GFP cells and Hoechst positive cells.

**Supplementary Figure 3:**
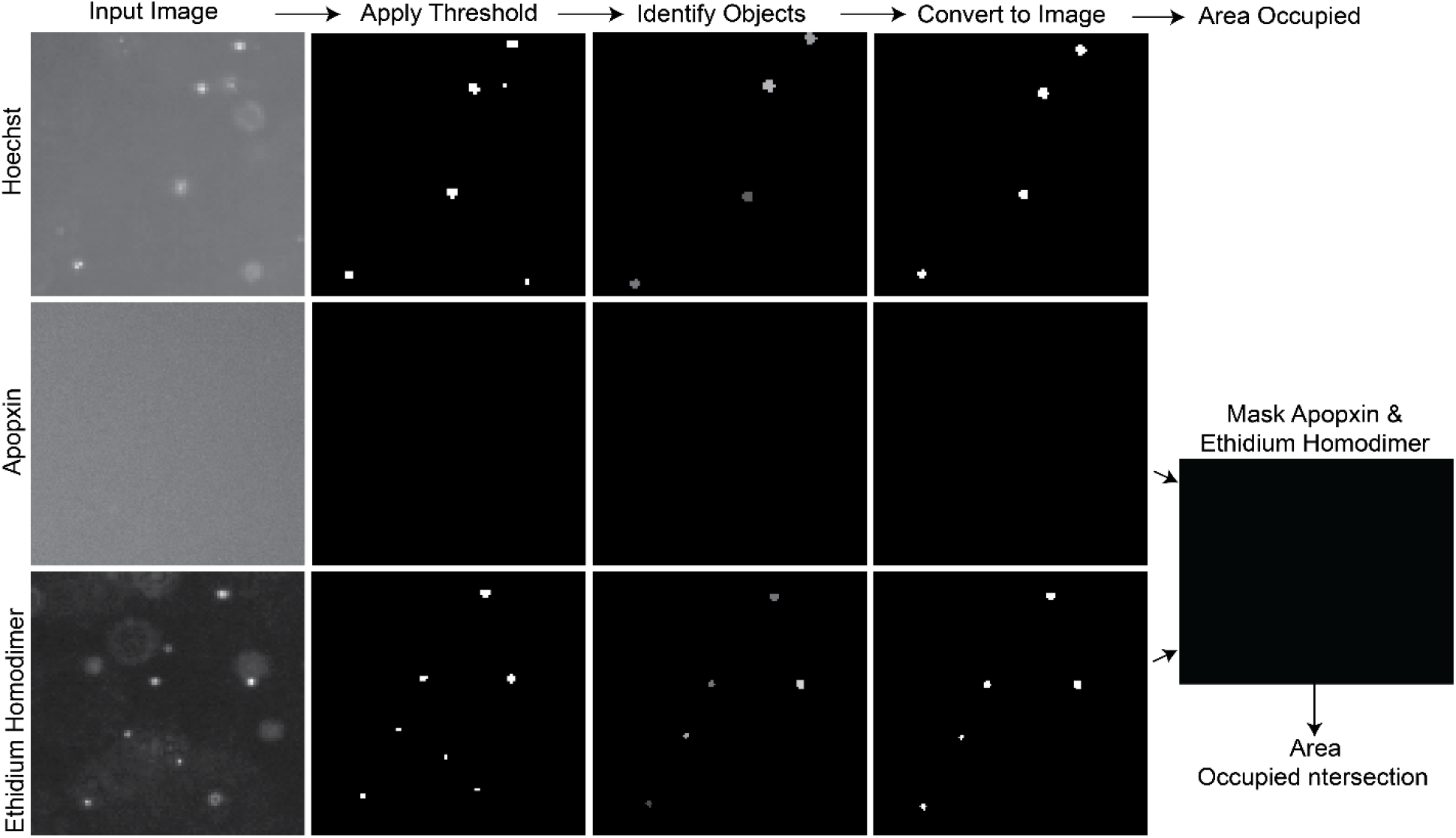
Cell profiler pipeline for image analysis: Image analysis using cell profiler involved thresholding, followed by segmentation to identify pixels to identify the total image area occupied for each channel. The intersection between EtHd and Apopxin was used to exclude pixels positive for both markers from being counted twice.

**Supplementary Figure 4:**
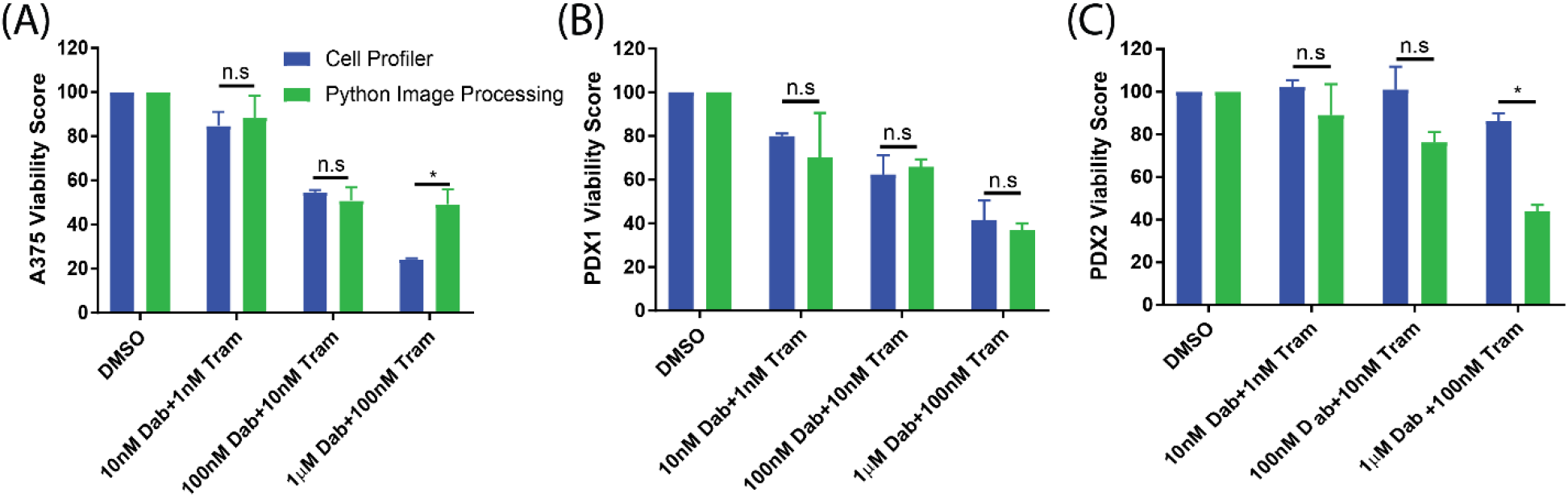
Comparison of results analyzed using Cell Profiler to analysis by python image analysis. Data from experiments using A375 **(A)** M481 **(B)** and M498 **(C)** cells treated with RMIC were analysis using both image analysis pipelines. Data shown are three independent experimental repeats with results presented as average ± SEM. (* p value < 0.05).

## References

1. Holohan C, Van Schaeybroeck S, Longley DB, Johnston PG: Cancer drug resistance: an evolving paradigm. Nature Reviews Cancer 2013, 13:714.

2. Maman S, Witz IP: A history of exploring cancer in context. Nature Reviews Cancer 2018, 18(6):359–376.

3. Poltavets V, Kochetkova M, Pitson SM, Samuel MS: The Role of the Extracellular Matrix and Its Molecular and Cellular Regulators in Cancer Cell Plasticity. Frontiers in oncology 2018, 8:431–431.

4. Leight JL, Drain AP, Weaver VM: Extracellular Matrix Remodeling and Stiffening Modulate Tumor Phenotype and Treatment Response. Annual Review of Cancer Biology 2017, 1(1):313–334.

5. Paszek MJ, Zahir N, Johnson KR, Lakins JN, Rozenberg GI, Gefen A, Reinhart-King CA, Margulies SS, Dembo M, Boettiger D et al: Tensional homeostasis and the malignant phenotype. Cancer Cell 2005, 8(3):241–254.

6. Wells RG: The role of matrix stiffness in regulating cell behavior. Hepatology 2008, 47(4):1394–1400.

7. Edmondson R, Broglie JJ, Adcock AF, Yang L: Three-Dimensional Cell Culture Systems and Their Applications in Drug Discovery and Cell-Based Biosensors. Assay and Drug Development Technologies 2014, 12(4):207–218.

8. LaBarbera DV, Reid BG, Yoo BH: The multicellular tumor spheroid model for high-throughput cancer drug discovery. Expert Opinion on Drug Discovery 2012, 7(9):819–830.

9. Santo VE, Rebelo SP, Estrada MF, Alves PM, Boghaert E, Brito C: Drug screening in 3D in vitro tumor models: overcoming current pitfalls of efficacy read-outs. Biotechnology Journal 2017, 12(1):1600505-n/a.

10. Su Y, Wei W, Robert L, Xue M, Tsoi J, Garcia-Diaz A, Homet Moreno B, Kim J, Ng RH, Lee JW et al: Single-cell analysis resolves the cell state transition and signaling dynamics associated with melanoma drug-induced resistance. Proceedings of the National Academy of Sciences 2017, 114(52):13679–13684.

11. Shaffer SM, Dunagin MC, Torborg SR, Torre EA, Emert B, Krepler C, Beqiri M, Sproesser K, Brafford PA, Xiao M et al: Rare cell variability and drug-induced reprogramming as a mode of cancer drug resistance. Nature 2017, 546:431.

12. Tirosh I, Izar B, Prakadan SM, Wadsworth MH, Treacy D, Trombetta JJ, Rotem A, Rodman C, Lian C, Murphy G et al: Dissecting the multicellular ecosystem of metastatic melanoma by single-cell RNA-seq. Science (New York, NY) 2016, 352(6282):189–196.

13. Antoine EE, Vlachos PP, Rylander MN: Review of collagen I hydrogels for bioengineered tissue microenvironments: characterization of mechanics, structure, and transport. Tissue engineering Part B, Reviews 2014, 20(6):683–696.

14. Wolf K, te Lindert M, Krause M, Alexander S, te Riet J, Willis AL, Hoffman RM, Figdor CG, Weiss SJ, Friedl P: Physical limits of cell migration: Control by ECM space and nuclear deformation and tuning by proteolysis and traction force. The Journal of Cell Biology 2013, 201(7):1069–1084.

15. Friedl P, Wolf K: Plasticity of cell migration: a multiscale tuning model. The Journal of Cell Biology 2010, 188(1):11–19.

16. Bendris N, Williams KC, Reis CR, Welf ES, Chen P-H, Lemmers B, Hahne M, Leong HS, Schmid SL: SNX9 promotes metastasis by enhancing cancer cell invasion via differential regulation of RhoGTPases. Molecular Biology of the Cell 2016, 27(9):1409–1419.

17. Welf Erik S, Driscoll Meghan K, Dean Kevin M, Schäfer C, Chu J, Davidson Michael W, Lin Michael Z, Danuser G, Fiolka R: Quantitative Multiscale Cell Imaging in Controlled 3D Microenvironments. Developmental Cell 2016, 36(4):462–475.

18. Kim BJ, Zhao S, Bunaciu RP, Yen A, Wu M: A 3D in situ cell counter reveals that breast tumor cell (MDA-MB-231) proliferation rate is reduced by the collagen matrix density. Biotechnology Progress 2015, 31(4):990–996.

19. Lv D, Yu S-C, Ping Y-F, Wu H, Zhao X, Zhang H, Cui Y, Chen B, Zhang X, Dai J et al: A three-dimensional collagen scaffold cell culture system for screening anti-glioma therapeutics. Oncotarget 2016, 7(35):56904–56914.

20. Long GV, Hauschild A, Santinami M, Atkinson V, Mandalà M, Chiarion-Sileni V, Larkin J, Nyakas M, Dutriaux C, Haydon A et al: Adjuvant Dabrafenib plus Trametinib in Stage III BRAF-Mutated Melanoma. New England Journal of Medicine 2017, 377(19):1813-1823.

21. Frantz C, Stewart KM, Weaver VM: The extracellular matrix at a glance. Journal of Cell Science 2010, 123(24):4195–4200.

22. Theocharis AD, Skandalis SS, Gialeli C, Karamanos NK: Extracellular matrix structure. Advanced Drug Delivery Reviews 2016, 97:4–27.

23. Bosman FT, Stamenkovic I: Preface to extracellular matrix and disease. The Journal of Pathology 2003, 200(4):421–422.

24. Mouw JK, Ou G, Weaver VM: Extracellular matrix assembly: a multiscale deconstruction. Nature Reviews Molecular Cell Biology 2014, 15:771.

25. Quintana E, Shackleton M, Sabel MS, Fullen DR, Johnson TM, Morrison SJ: Efficient tumour formation by single human melanoma cells. Nature 2008, 456:593.

26. Han YL, Ronceray P, Xu G, Malandrino A, Kamm RD, Lenz M, Broedersz CP, Guo M: Cell contraction induces long-ranged stress stiffening in the extracellular matrix. Proceedings of the National Academy of Sciences 2018.

27. Hirata E, Girotti Maria R, Viros A, Hooper S, Spencer-Dene B, Matsuda M, Larkin J, Marais R, Sahai E: Intravital Imaging Reveals How BRAF Inhibition Generates Drug-Tolerant Microenvironments with High Integrin β1/FAK Signaling. Cancer Cell 2015, 27(4):574–588.

28. Hirata E, Sahai E: Tumor Microenvironment and Differential Responses to Therapy. Cold Spring Harbor Perspectives in Medicine 2017.

29. Antoni D, Burckel H, Josset E, Noel G: Three-Dimensional Cell Culture: A Breakthrough in Vivo. International Journal of Molecular Sciences 2015, 16(3):5517-5527.

30. Friedrich J, Seidel C, Ebner R, Kunz-Schughart LA: Spheroid-based drug screen: considerations and practical approach. Nature Protocols 2009, 4:309.

31. Timmins NE, Nielsen LK: Generation of Multicellular Tumor Spheroids by the Hanging-Drop Method. In: Tissue Engineering. edn. edited by Hauser H, Fussenegger M. Totowa, NJ: Humana Press; 2007: 141–151.

32. Foty R: A simple hanging drop cell culture protocol for generation of 3D spheroids. Journal of visualized experiments : JoVE 2011(51):2720.

33. Mathew B, Schmitz A, Muñoz-Descalzo S, Ansari N, Pampaloni F, Stelzer EHK, Fischer SC: Robust and automated three-dimensional segmentation of densely packed cell nuclei in different biological specimens with Lines-of-Sight decomposition. BMC Bioinformatics 2015, 16:187.

34. Macarron R, Banks MN, Bojanic D, Burns DJ, Cirovic DA, Garyantes T, Green DVS, Hertzberg RP, Janzen WP, Paslay JW et al: Impact of high-throughput screening in biomedical research. Nat Rev Drug Discov 2011, 10(3):188–195.

35. Workman P, Draetta GF, Schellens JHM, Bernards R: How Much Longer Will We Put Up With $100,000 Cancer Drugs? Cell 2017, 168(4):579–583.

36. Hongisto V, Jernström S, Fey V, Mpindi J-P, Kleivi Sahlberg K, Kallioniemi O, Perälä M: High-Throughput 3D Screening Reveals Differences in Drug Sensitivities between Culture Models of JIMT1 Breast Cancer Cells. PLOS ONE 2013, 8(10):e77232.

37. Kenny HA, Krausz T, Yamada SD, Lengyel E: Use of a novel 3D culture model to elucidate the role of mesothelial cells, fibroblasts and extra-cellular matrices on adhesion and invasion of ovarian cancer cells to the omentum. International Journal of Cancer 2007, 121(7):1463–1472.

38. Kenny HA, Lal-Nag M, White EA, Shen M, Chiang C-Y, Mitra AK, Zhang Y, Curtis M, Schryver EM, Bettis S et al: Quantitative high throughput screening using a primary human three-dimensional organotypic culture predicts in vivo efficacy. Nature Communications 2015, 6:6220.

39. Packard RRS, Baek KI, Beebe T, Jen N, Ding Y, Shi F, Fei P, Kang BJ, Chen P-H, Gau J et al: Automated Segmentation of Light-Sheet Fluorescent Imaging to Characterize Experimental Doxorubicin-Induced Cardiac Injury and Repair. Scientific Reports 2017, 7(1):8603.

40. Huisken J, Stainier DYR: Even fluorescence excitation by multidirectional selective plane illumination microscopy (mSPIM). Optics Letter 2007, 32(17):2608–2610.

